# Cortical motor activity modulates respiration and reduces apnoea in neonates

**DOI:** 10.1101/2025.04.03.646882

**Authors:** Coen S. Zandvoort, Fatima Usman, Shellie Robinson, Odunayo Fatunla, Eleri Adams, Kyle T.S. Pattinson, Simon F. Farmer, Caroline Hartley

**Affiliations:** Department of Paediatrics, University of Oxford, Oxford, United Kingdom; Newborn Care Unit, John Radcliffe Hospital, Oxford University Hospitals NHS Foundation Trust, Oxford, United Kingdom; Nuffield Department of Clinical Neurosciences, University of Oxford, Oxford, UK; Nuffield Department of Anaesthetics, Oxford University Hospitals NHS Foundation Trust, Oxford, UK; Department of Neurology, National Hospital for Neurology and Neurosurgery, London, United Kingdom; Department of Clinical and Human Neuroscience, UCL Institute of Neurology, London, United Kingdom

## Abstract

Respiration is governed by a widespread network of cortical and subcortical structures. This complex communication between the brain and lungs is altered in pathological conditions. Apnoea – the cessation of respiration – is a common condition in infants, particularly those born prematurely. Apnoea in infants is believed to relate to immaturity of brainstem respiratory centres; involvement of the cortex in respiration in infants has yet to be explored. We investigated if there was any evidence for cortical coupling with respiration in newborn humans and whether it relates to apnoea. Using simultaneous electroencephalography (EEG) and impedance pneumography we investigated interactions between cortical and respiratory activity (known as cortico-respiratory coupling) using phase-amplitude coupling. We show that cortico-respiratory coupling is present in premature and term newborns (104 recordings from 68 infants; 34.5 ± 2.6 weeks post-menstrual age), identifying an interplay between breathing phase and EEG amplitude. We further shed light on the biological meaning by revealing that the strongest coupling occurs during inspiration and that cortical activity precedes respiration, with coupling strongest over frontocentral regions. Whilst our study was limited in spatial resolution, and determining causality is challenging, we believe these findings support the notion that the cortico-respiratory coupling observed here constitutes communication between cortical motor areas and lung effectors. Moreover, we show that cortico-respiratory coupling is negatively correlated with the rate of apnoea, revealing novel insight into this common and potentially life-threatening neonatal pathology.

## Introduction

Apnoea – the cessation of respiration – is a potentially life-threatening event^1^, affecting more than 50% of preterm infants and some term infants^2^. Apnoeic episodes can result in physiological instability including oxygen desaturation and bradycardia^1^, are associated with suppression of brain activity^3^, and have been linked to poorer neurodevelopmental outcomes^4^. Apnoea of prematurity is caused by immaturity of the lungs, chest wall, and nervous system^5^. Immaturity of the brainstem plays a significant role in apnoea^1^; however, the relationship with other respiratory-linked brain regions is less clear. In adults with congenital central hypoventilation syndrome^6^ or obstructive sleep apnoea^7^, a respiratory-linked increase in cortical motor activity suggests that the motor cortex plays an important role in maintaining autonomous respiration, with the authors postulating that cortico-respiratory drive whilst participants are awake may prevent the hypoventilation/apnoea observed in these patients whilst they are asleep.

Recently, it was proposed that communication between the cortex and lungs, known as cortico-respiratory coupling, can be identified and quantified through phase-amplitude coupling^8,9^. This functional coupling arises when the amplitude of cortical activity is modulated by the respiration phase, or *vice versa*. Phase-amplitude coupling is typically quantified using non-invasive recordings capturing respiratory and neural activity (e.g., magneto- or electroencephalography [EEG]). Studies have demonstrated that the respiratory phase at low frequencies (<1 Hz) relates to higher-frequency oscillations in the cortex^9,10^. Phase-amplitude coupling therefore allows us to quantify the interplay between respiratory and cortical activity. In adults, cortico-respiratory coupling can be observed in a wide network of subcortical and cortical regions including motor areas^9^. Moreover, cortico-respiratory coupling is aberrated in pathological conditions^11^.

Here, using phase-amplitude coupling, we investigated whether cortical areas are coupled with respiration in infants (Figure 1). We hypothesised that cortico-respiratory coupling occurs in newborns and that the strength of cortico-respiratory coupling is negatively associated with apnoea rate (in line with the suggestions made from studies of adults with congenital central hypoventilation syndrome^6^ and obstructive sleep apnoea^7^). To this end, we first examined whether cortico-respiratory coupling exists in both premature and term infants. To investigate if the coupling might relate to central motor drive, we further studied the spatial localisation, timing throughout the respiratory cycle and coupling directionality. Finally, we examined the relationship between cortico-respiratory coupling and apnoea rate.

**Figure 1.**
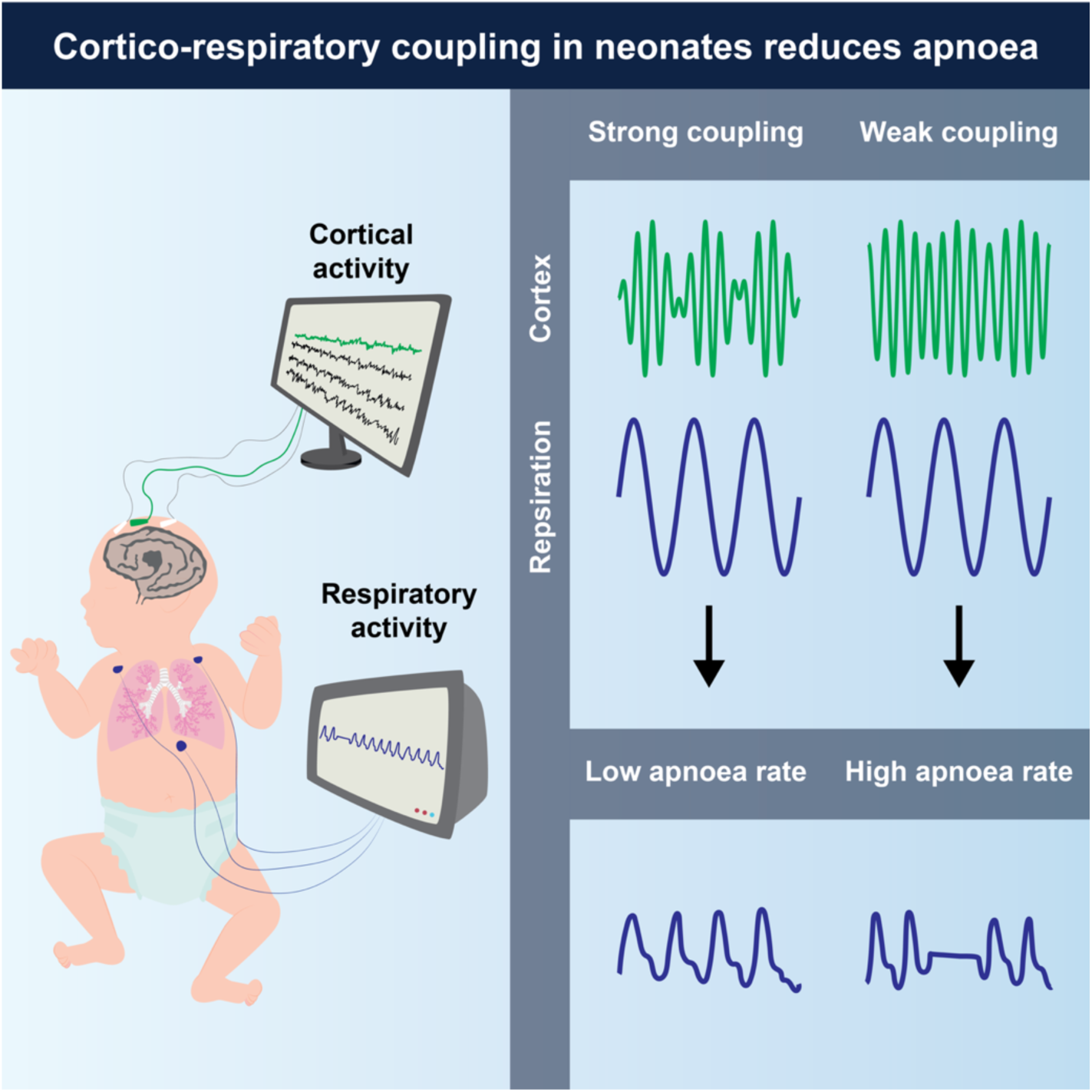
Schematic of the data acquisition setup (left panel) and main findings (right panel). Each recording consisted of cortical activity (using electroencephalography) and respiratory activity (using impedance pneumography).

## Results and Discussion

### Respiratory and cortical activity exhibit phase-amplitude coupling in infants

We analysed simultaneous 8-channel resting-state EEG (to measure cortical activity) and impedance pneumography (IP; to measure respiration) in 68 infants (28-42 weeks postmenstrual age [PMA] at time of recording) on 104 occasions. We first assessed whether and how EEG and IP are interrelated using cross-frequency coupling^9,12,13^. In line with findings in adults^9^, we expected that the amplitude of oscillatory brain activity is modulated as a function of the respiratory cycle. That is, we expected to detect an EEG component that systematically varied with timing within the respiratory cycle. To this end, we identified individual breaths (and thus respiratory phase) from the IP signal^14^. Through averaging the EEG (time-locked to the IP signal) across breaths, we observed that EEG amplitude was modulated in the delta- (0.5-4 Hz) and theta-band (4-8 Hz) frequencies depending on the respiratory phase (Figure 2A-B). This analysis can be seen as a first indication that the EEG amplitude in specific frequency ranges changes systematically with the respiratory phase.

**Figure 2.**
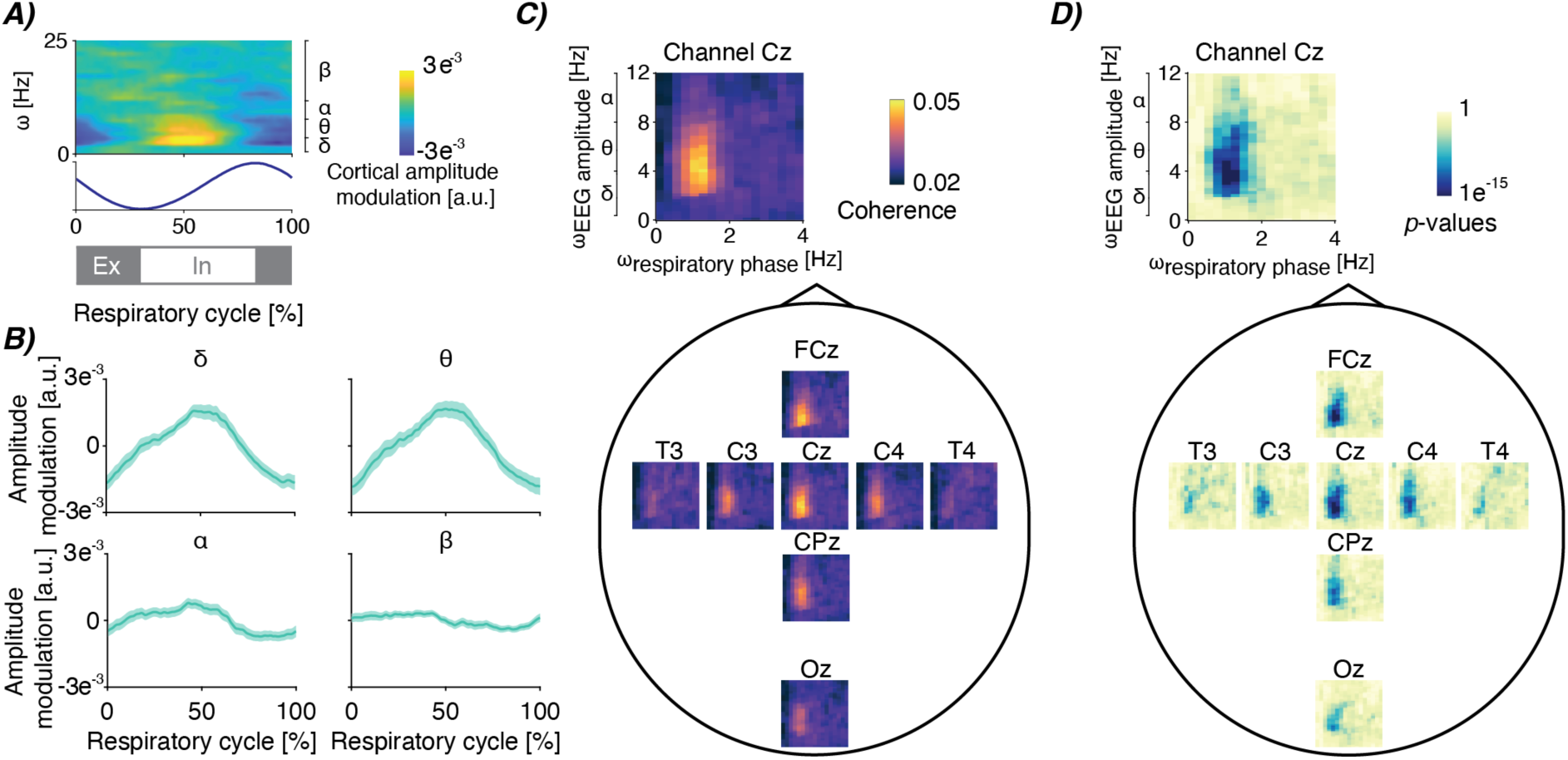
Cortico-respiratory coupling in infants. **A)** Electroencephalographic (EEG) amplitude modulation depending on the respiratory cycle and frequency. EEG amplitude is averaged over breaths, recordings and channels FCz and Cz. The blue graph below is the impedance pneumographic signal time-locked from breath to breath and averaged over breaths and recordings. Grey and white bars below indicate expiration (Ex; in grey) and inspiration (In; in white). **B)** The time-resolved amplitude modulations averaged within four frequency bands (delta: 0.5-4 Hz, theta: 4-8 Hz, alpha: 8-13 Hz, beta: 13-25 Hz), focusing on channels FCz and Cz again. Shaded light-green area is the standard error across recordings after pooling over channels. **C)** Spatial map of coherence-based phase-amplitude coupling (PAC) between the respiratory phase (ω_respiratory phase_) and EEG amplitude (ω_EEG amplitude_) – see Figure S1 for the pan-spectral PAC estimates. Spectra are the mean over all recordings. **D)** Spatial map for the corresponding (uncorrected) statistical significance relative to surrogate PAC obtained through epoch shuffling (reported on a logarithmic scale). For all panels, data included is from 68 infants (28-42 weeks postmenstrual age [PMA] at time of recording) on 104 recording occasions. See Table 1 for further clinical and demographic characteristics.

**Table 1.** Infant demographics. Reported values are number of babies or recordings, or mean and standard deviation [SD]/median and interquartile range [IQR] – as indicated. Values are reported over recordings for ages (postmenstrual, gestational, and postnatal), respiratory distress syndrome, persistent pulmonary hypertension, infection and respiratory support. Birthweight, sex, mode of delivery, and Apgar scores are provided over infants.

|  |  |
| --- | --- |
| Postmenstrual age at recording (weeks, mean±SD) | 34.5 ± 2.6 |
| Gestational age at birth (weeks, mean±SD) | 31.1 ± 3.4 |
| Postnatal age at recording (weeks, mean±SD) | 3.4 ± 2.2 |

**Table 1. Infant demographics.** Reported values are number of babies or recordings, or mean and standard deviation [SD]/median and interquartile range [IQR] – as indicated.
|  |  |
| --- | --- |
| **Birthweight (g, mean±SD)** | 1,726 ± 816 |
| **Sex** |  |
| Females (n) | 27 |
| Males (n) | 47 |
| **Mode of delivery** |  |
| Normal vaginal delivery (n) | 16 |
| Vaginal assisted (ventouse/forceps/kiwi) (n) | 4 |
| Elective Caesarean section (n) | 7 |
| Emergency Caesarean section (n) | 47 |
| **Apgar scores** |  |
| Apgar at 1 minute (median [IQR]) | 8 [3] |
| Apgar at 5 minutes (median [IQR]) | 10 [2] |
| Apgar at 10 minutes (median [IQR]) | 10 [0] |
| **Respiratory distress syndrome** |  |
| Ongoing condition (n) | 39 |
| Past treated (n) | 48 |
| No (n) | 36 |
| **Persistent pulmonary hypertension** |  |
| Past treated (n) | 5 |
| No (n) | 118 |
| **Infection** |  |
| None (n) | 48 |
| Treated (n) | 75 |
| **Respiratory support** |  |
| High flow therapy (n) | 25 |
| Low flow therapy (n) | 16 |
| Spontaneous ventilation (n) | 82 |
| **Caffeine received on the day of the EEG recording** |  |
| Yes (n) | 59 |
| No (n) | 64 |

We then statistically confirmed this association between EEG and respiration using phase-amplitude coupling (PAC). Using cross-frequency coherence, we observed significant PAC between the phase at the mean respiratory frequency and the EEG amplitude in delta and theta frequency bands (Figure 2C-D). PAC was strongest at central electrodes (channels FCz and Cz; Figure 2C-D; *p*<0.001 false discovery rate [FDR] corrected). In line with the results presented in Figure 2A, we did not find significant PAC at higher EEG frequencies such as the alpha (8-13 Hz) and beta (13-25 Hz) frequency bands (Figure S1).

These findings confirm that cortico-respiratory coupling occurs in infants. Whilst we are the first to show that cortico-respiratory coupling already exists in early life, adult studies have revealed widespread brain-respiratory networks covering both cortical and subcortical areas, as well as pan-spectral components of oscillatory brain activity (2-150 Hz)^9^. These networks have been linked to a diverse range of perceptual, cognitive and motor processes^8,15–17^. In these studies, respiratory rhythm has been found to relate to task performance (e.g., better performances are found during a certain respiratory phase) as well as neural oscillations in the brain^11^. For example, during an isometric motor task, the strength of cortico-muscular communication at beta-band frequencies cyclically changes with the respiratory phase^16^. Moreover, the probability of initiating goal-directed movements relates to cortical motor activity (measured as the readiness potential amplitude) and respiratory phase^17^. These relationships between task performance, respiration and cortical activity are well-established and replicated in a variety of animal and human adult studies, demonstrating the importance of cortico-respiratory coupling.

### Cortico-respiratory coupling is strongest during inspiration and may be driven by cortical activity

To further shed light on the functional meaning of cortico-respiratory coupling in neonates, we next examined the timing and directionality of the coupling. To study whether coupling strength fluctuates throughout the respiratory cycle, we focused on the EEG channels with strongest coupling (FCz and Cz), and frequencies where statistically significant coupling occurs (Figure 2D; *p*<0.001 FDR-corrected). We confirmed that PAC changes in a time-dependent manner, observing the strongest PAC during the inspiratory phase (Figure 3A-B). Such within-breath modulation is evident for both delta- and theta-frequencies (Figure 3A-B). This indicates that the strength of cortico-respiratory coupling changes throughout the respiratory cycle. During the inspiratory phase, neural activation from the central nervous system is required to drive respiratory motoneurons causing contraction of respiratory muscles. Given the phasic PAC modulation as seen in Figure 3A-B, this suggests that cortical areas centred around the vertex may play an important role in the neural activation of respiratory muscles.

**Figure 3.**
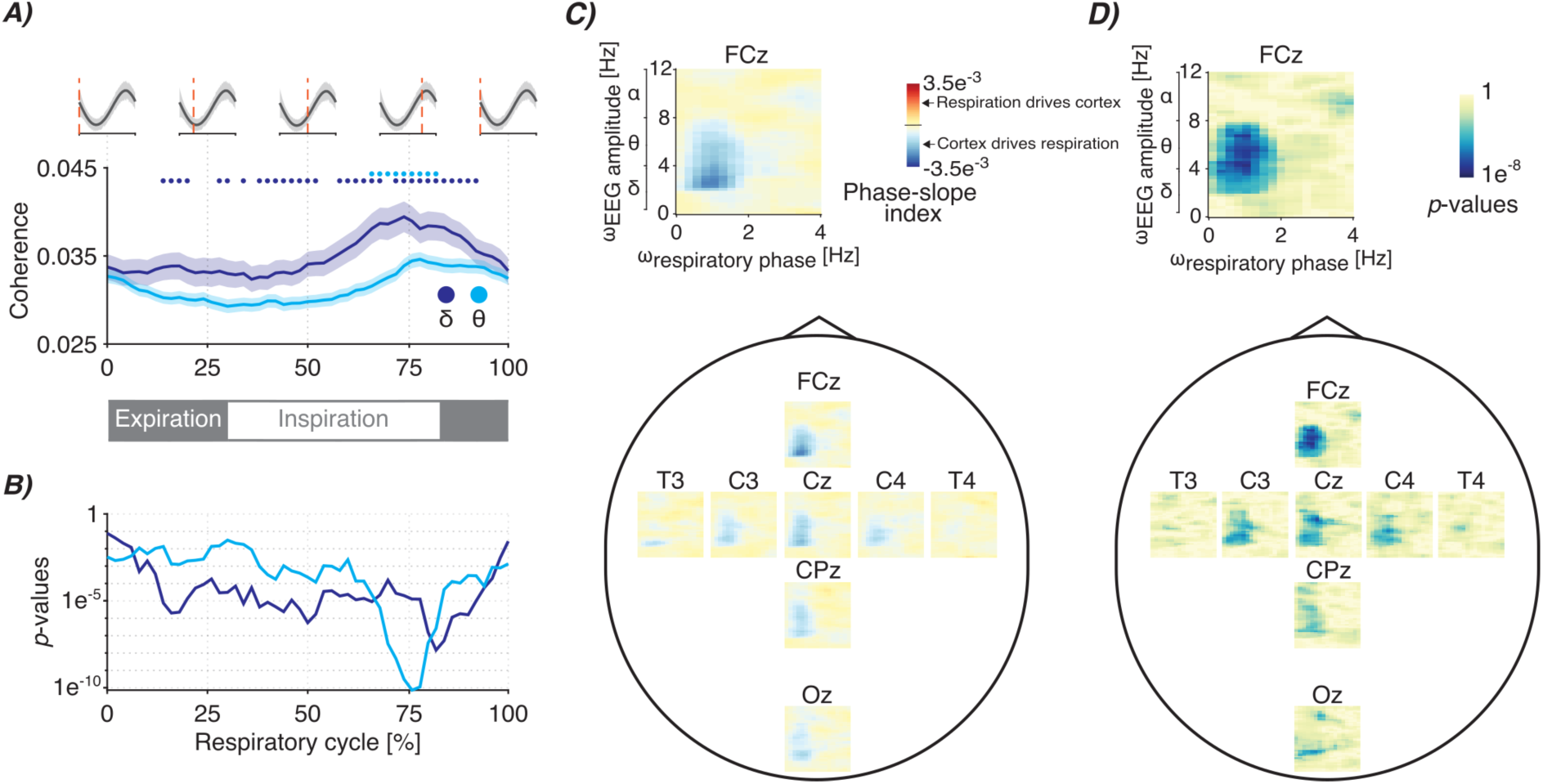
Timing and directionality of the cortico-respiratory coupling. **A)** Phase-amplitude coupling modulation (estimated as cross-frequency coherence) throughout the respiratory cycle. Graphs are colour-coded for the delta- and theta-frequency bands (dark- and light-blue colours respectively), which are the EEG’s primary frequency bands for which we observed statistically significant PAC (Figure 2C-D). Shaded regions indicate standard error over recordings. PAC was estimated for EEG channels FCz and Cz. Grey waves at the top and grey bar at the bottom indicate the phase of the respiratory cycle. Coloured dots indicate points that are statistically significant (alpha level was set to 0.001 with false discovery rate correction). **B)** Corresponding (uncorrected) statistical significance analogous to the coloured dots at the top of panel A. **C)** Spatial map of the cross-frequency phase-slope index between respiratory phase (ω_respiratory phase_) and EEG amplitude (ω_EEG amplitude_) (see Figure S2 for the pan-spectral estimates) and **D)** its (uncorrected) statistical significance (reported on a logarithmic scale). The spectra show the mean over all recordings. For all panels, data included is from 68 infants (28-42 weeks postmenstrual age [PMA] at time of recording) on 104 recording occasions. See Table 1 for further clinical and demographic characteristics.

Standard coherence (i.e., our PAC measure) only quantifies coupling in a bidirectional manner and cannot directly show any directionality, meaning that it is not possible to distinguish whether cortical activity precedes or follows respiratory activity. To investigate whether cortical activity precedes respiration or *vice versa*, we employed a cross-frequency version of the phase-slope index^13^. This revealed that delta- and theta-band EEG amplitude leads respiratory activity (Figure 3C). We observed the strongest phase-slope index around central areas, in particular for channel FCz (Figure 3D; *p*<0.05 FDR-corrected). Moreover, in line with the PAC analysis (Figure 2), phase-slope index was not statistically significant for alpha- and beta-band frequencies (Figure S2). However, caution is needed in the interpretation of these results as signal processing techniques such as the phase-slope index estimate directionality but do not confirm causality. Rather, they show a statistical relationship which can be influenced by a multitude of factors (e.g., signal-to-noise ratio and preprocessing steps). Nevertheless, the results suggest that cortical activity may precede respiration in newborns. Future work is needed to confirm this association by, for example, employing other metrics to estimate directionality, such as the time-lagged cross-correlation and Granger causality and through direct mechanistic studies.

Considering the cortico-respiratory coupling’s spatial location, its timing in the respiratory cycle and its directionality of action, combined with previous work in adults and animal models, leads us to suggest that this coupling observed may represent the respiratory drive from higher-order motor centres, primarily encompassing the cortical motor areas. It is important to note that the number of electrodes in our montage (with only 8 recording electrodes) is limited, and so source localisation was not possible; higher-density recordings are warranted to confirm whether the motor cortex plays a role. Structurally, the motor cortex connects to respiratory muscles both directly and indirectly^18^. This allows for functional control over respiratory muscles (comprising diaphragm and intercostal muscles) via the corticospinal^19^ and reticulospinal tracts^20^. In cat preparations, transection of the ventrolateral part of the spinal cord (which essentially blocks the brainstem input to respiratory motoneurons) does not prevent cortical stimulation from eliciting responses in the respiratory intercostal muscle motoneurons, signifying the cortico-spinal drive^20^. Such motor-evoked potentials can also be elicited by stimulating the cortical motor areas of humans using transcranial magnetic stimulation^21,22^.

Whilst we believe that it is possible that our coupling reflects cortical motor drive, there are several limitations to consider and further mechanistic studies are warranted. One might also argue that cortico-respiratory coupling could reflect changes in blood flow or oxygen saturation which induces ‘coupling’, where blood flow/oxygen saturation acts as a confounding factor essentially modulating respiratory and brain activity in a cycle-dependent manner. It is, however, unlikely that this is the case for our results given the focal spatial localisation around central electrodes and narrow spectral bandwidth comprising EEG frequencies between 1-8 Hz. Moreover, the amplitude modulation of blood flow reveals a different time-frequency representation over the respiratory cycle (Figure S3) when compared to cortical activity (Figure 2A).

To further our understanding of whether the coupling in infants relates directly to central motor drive, future work should examine potential coupling in the cortico-muscular domain. Electrical activity of the diaphragm (Edi) in infants can be recorded using an Edi catheter^23^, as is used in ventilators with neurally adjusted ventilatory assist^24^. Correlating this electromyographic (EMG) activity from the diaphragm with brain activity (e.g., EEG) will enable cortico-muscular assessments that can quantify whether oscillatory components in the EEG and diaphragmatic EMG are phase-locked. It is of particular interest whether linear coherence can be observed in the frequency range of 1-8 Hz, which is expected from our PAC findings presented in Figure 2. Interestingly, such coherence analysis has been conducted at the muscular level in rodents. Invasively recording activity from the diaphragm and intercostal muscles revealed strong inter-muscular coherence in the lower frequencies between 1 and 8 Hz^25^. Future work should also establish to what extent cortical motor activity is related to other respiratory functionalities such as altering respiratory rate. In the current investigation, we focused on how cortical amplitude directly related to the respiratory phase but phase-amplitude coupling of cortical activity has also been revealed to play a role in adjusting respiratory rate^26^ (i.e., respiratory rate rather than individual breaths).

### Cortico-respiratory coupling relates to apnoea in infants

Having identified significant cortico-respiratory coupling in infants, we then tested our hypothesis that the strength of the cortico-respiratory coupling is negatively associated with apnoea. Apnoea of prematurity can lead to periods of physiological instability when apnoea is accompanied by bradycardia and desaturation^5^. These episodes may have long-term neurological effects^4^ and, hence, it is important to better understand the pathophysiology. In addition to pulmonary immaturity, apnoea of prematurity has been primarily attributed to immaturity and weakened neural drive from the respiratory centres in the brainstem. These respiratory centres are principally located in the medulla oblongata and pons within the brainstem, which ensure the periodical activation of lung muscles to enable respiration. However, as our results suggest that further central input to respiration may be provided by higher-order brain areas such as the cortical motor areas, investigating whether there is an association between cortico-respiratory coupling and apnoea in infants is warranted.

To test our hypothesis, apnoea rate was estimated over a time window of up to 24 hours (infants have their vital signs continuously monitored at the neonatal care unit), whereas cortico-respiratory coupling was assessed on the same day from EEG recordings with an average duration of 1.7 ± 0.6 hours. Choosing different time windows ensured that any relationship between apnoea rate and cortico-respiratory coupling was not the mere result of the number of breaths included in the analysis. For instance, when the same respiratory signal is used in both analyses, a higher apnoea rate would mean that fewer windows are used for the PAC estimation which could result in an increased magnitude. We selected PAC pooled over EEG channels Cz and FCz since their coupling magnitudes were the highest.

We found support for our hypothesis that apnoea and cortico-respiratory-coupling are related. PAC was negatively associated with apnoea rate (*t*(98):-2.37, *p*:0.02, *ρ*:-0.23; Figure 4A; this relationship was also identified when focusing on individual channels [Figure S4] and preterm infants [≤36 weeks, Figure S5]). This suggests that cortico-respiratory coupling may relate to the likelihood of infant’s experiencing apnoeas. As caffeine is given to infants for the treatment of apnoea of prematurity, we also conducted a stratified analysis according to whether infants were receiving caffeine at the time of the EEG recording or not (Figure S6). Conducting a stratified analysis substantially reduces sample size, nevertheless, there was a strong trend towards a relationship between apnoea rate and PAC in the infants receiving caffeine, who are the infants most likely to experience apnoea (Figure S6B).

**Figure 4.**
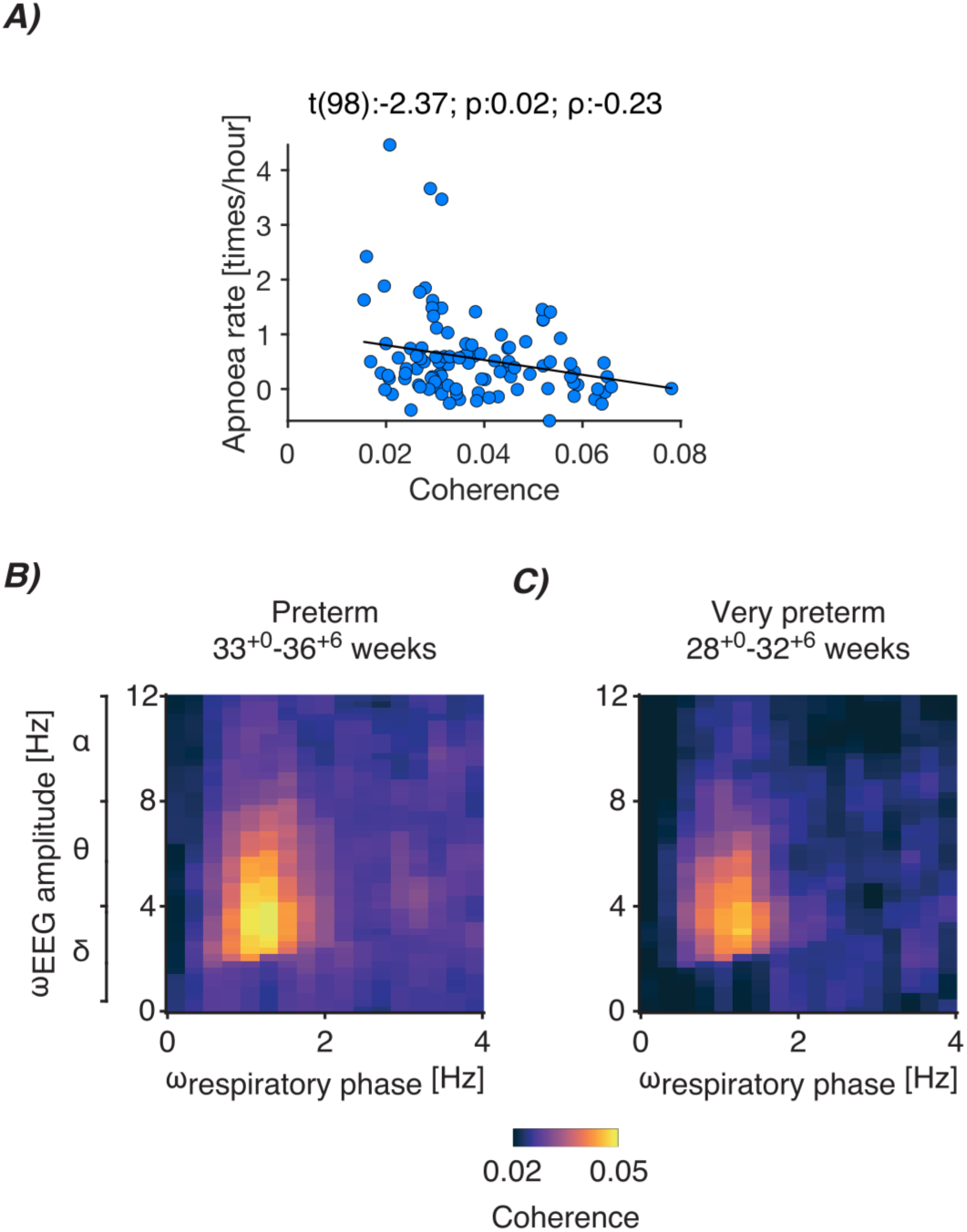
Cortico-respiratory coupling dependency on apnoea rate and postmenstrual age. **A)** Relationship between apnoea rate and statistically significant phase-amplitude coupling (PAC; defined as coherence encompassing delta- and theta-band frequencies). Each blue dot is the data of an individual recording (data of 68 infants were included from 104 test occasions). Black linear graph is the best fit of the linear mixed-effects model (fixed factors: coupling averaged over statistically significant samples from channels FCz and Cz [see Figure S4 for channel-specific relationships], data length to determine apnoea rate, mode of ventilation, and postmenstrual age; random factor: infant). Panel title contains the test statistic of the predictor’s regression slope (t), its significance (p), and partial correlation coefficient (ρ). See Figures S5-S7 for data according to PMA and whether or not the infant received caffeine at the time of recording. **B-C)** PAC spectra between the respiratory phase (ω_respiratory phase_) and EEG amplitude (ω_EEG amplitude_) for **B)** preterm (58 recordings; 33.0-36.86 weeks), and **C)** very preterm (28 recordings; 28.0-32.86 weeks) infants. Both spectra are averaged over recordings and EEG channels FCz and Cz.

Overall, the negative association between apnoea rate and PAC provides further support that our measures of cortico-respiratory coupling (Figures 2-3) are related to central motor drive. It is plausible that in infants cortical motor activity assists the brainstem’s central pattern generators (CPGs) to establish increased control over the respiratory muscles, which in our analysis, is reflected as an association with apnoea rate. The motor cortex can exert coordination over the CPGs through its downstream projections to the thalamus^27^, which, in turn is connected to the preBötzinger complex (the structure that primarily hosts the CPGs^28^). Nevertheless, it is important to recognise that a limitation of this analysis is that correlation does not imply causation, and future mechanistic studies are required to determine whether and how cortico-respiratory coupling plays a role in reducing apnoea in infants.

In theory, the strength of cortico-respiratory coupling and its inverse relationship with apnoeas could reflect a shared relationship with PMA, where both coupling strength and apnoea rate change depending on PMA. The apnoea rate is expected to decrease with increased PMA across a broad age range^29^. It is also plausible that the strength of cortico-respiratory coupling (i.e., PAC magnitude) may change with PMA, with younger (i.e., preterm) infants potentially having weaker coupling. However, in our dataset, we observe statistically significant PAC in preterm infants (33^+0^-36^+6^ weeks^+days^; Figure 4B), and very preterm infants (28^+0^-32^+6^ weeks^+days^; Figure 4C). Moreover, PAC and apnoea rate were not significantly associated with PMA (Figure S7A-B, linear mixed-effects analysis). Similarly, PAC and apnoea rate were not significantly related to postnatal age (PNA; defined as the age in weeks from birth; Figure S7C-D). Whilst future studies should investigate the relationship between cortico-respiratory coupling and age across an even broader age range, the lack of significant associations here suggests that the relationship we observed between apnoea rate and cortico-respiratory coupling is not determined by age.

It is known that most apnoeas in infants occur during active sleep^6,30^ and delta-and theta-band frequencies in EEG are strongly related to sleep state^31^. A limitation of our study is that we did not record the sleep state of the infant. We speculate that cortico-respiratory coupling may change depending on sleep state, giving rise to more episodes of apnoea during certain sleep stages. Indeed, adult studies have suggested that cortical respiratory drive may occur predominantly during wakefulness^6,32^. Future work should study the effects of sleep stages on cortico-respiratory coupling to shed further light on the role of cortical motor drive for respiration in infants.

In summary, we demonstrate the existence of cortico-respiratory coupling in newborn infants. We believe that this coupling may be related to central motor drive due to its spatial location (predominantly over frontocentral EEG channels), timing (during inspiration) and directionality (with the cortex driving respiration). Consistent with our hypothesis, we found a significant relationship between apnoea rate and coupling strength, with infants with stronger cortico-respiratory coupling less likely to experience apnoea. The limitations of our study need to be considered, and in particular, directionality of the cortico-respiratory coupling, improved spatial localisation, and a direct mechanistic link between cortico-respiratory coupling and apnoea rate, should be investigated in future work. Nevertheless, our results have far-reaching implications for our understanding of apnoea of prematurity as they suggest that cortical motor areas play a pivotal role in apnoea in infants.

## Data and code availability

- Due to ethical restrictions, the raw data that support the findings of this study are available on reasonable request. Requests can be made directly to the corresponding author or directed to the Paediatric Neuroimaging Group (University of Oxford) through the institutional. Researchers will be required to agree to destroy the data within the time limits stipulated in the ethically approved study protocols. Upon agreement, data will be provided within 1 month.
- Summary-level data (before regression model adjustments of Figure 4), and channel- and recording-specific PAC spectra are provided at https://gitlab.com/paediatric_neuroimaging/cortico_respiratory_coupling.
- The codes for all analyses are available at: https://gitlab.com/paediatric_neuroimaging/cortico_respiratory_coupling. The algorithm for inter-breath interval detection is online at: https://gitlab.com/paediatric_neuroimaging/identify_ibi_from_ip.
- Any additional information required to reanalyse the data reported in this paper is available from the lead contact upon request.

## Acknowledgements

This work is funded by the Wellcome Trust and Royal Society through a Sir Henry Dale Fellowship awarded to Caroline Hartley (grant number: 213486/Z/18/Z). Fatima Usman is funded by the Commonwealth Scholarship Commission. Odunayo Fatunla is funded by the Clarendon Fund in partnership with the Lincoln College Kingsgate Graduate Scholarship. We would like to thank members of the Paediatric Neuroimaging Group (University of Oxford) for help with data collection, and the babies and their parents who took part in this research.

## Author contributions

**Conceptualization:** CSZ and CH; **Data curation:** FU, SR, OF, and CH; **Formal analysis:** CSZ; **Funding acquisition:** CH; **Investigation:** FU, SR, and OF; **Methodology:** CSZ; **Project administration:** CH; **Software:** CSZ; **Resources:** EA, KTSP and SFF; **Supervision:** CH and EA; **Visualization:** CSZ; **Writing – original draft:** CSZ; **Writing – review & editing:** all authors

## Material & Methods

### Participants and study design

This study was conducted at the Newborn Care Unit of the John Radcliffe Hospital, Oxford University Hospitals NHS Foundation Trust, Oxford, UK. The National Research Ethics Service approved the study (references: 19/LO/1085; 12/SC/0447). Eligible families were approached and given verbal and written details about the study. Parents provided written consent to take part in the study. The study conformed to the standards set by the Declaration of Helsinki and Good Clinical Practice.

Clinically stable infants aged between 28- and 42-weeks postmenstrual age (PMA) at the time of the test occasion were eligible to be included in the study. Infants were excluded if they had intraventricular haemorrhage grades III or IV, hypoxic ischaemic encephalopathy, congenital malformations, there was a known history of maternal substance misuse during pregnancy, or at the time of the study, they were receiving mechanical ventilation, opioid analgesics, or antibiotics (as a surrogate marker of infection). These conditions are known to alter respiratory and brain activity. We enrolled 74 infants who were recorded on 123 separate occasions (range: 1-8 recordings per infant). Fifty-one infants were included in one or two test occasions; 23 infants took part in the ongoing *Breathing and Brain Development* study where they are studied approximately once a week whilst in the Newborn Care Unit (included on 3±2 [median ± interquartile range] occasions). Demographic details of all infants can be found in Table 1. Parts of this dataset have been reported earlier in Zandvoort et al. ^33^ to address a different research question (this study investigated the development of sensory-evoked potentials, which were also recorded in these infants; it did not explore respiration).

### Data recordings

During each test occasion, approximately one to two hours of simultaneous electroencephalography (EEG) and vital signs (respiration, heart rate and oxygen saturation) were recorded. We continuously recorded the vital signs throughout their hospital stay of the infants who took part in the *Breathing and Brain Development* study.

Vital signs were recorded for each infant using Phillips IntelliVue MX800 or MX750 monitors. These data were continuously downloaded from the vital signs monitor using an electronic data capture software (iXtrend, ixitos, Germany). Both electrocardiograph (ECG; sampling frequency: 250 Hz) and impedance pneumography (IP; sampling frequency: 62.5 Hz) were recorded with three electrodes placed on the infant’s chest. Peripheral photoplethysmography was recorded at 125 Hz from a probe placed on the infant’s foot or hand.

EEG was recorded from DC to 400 Hz using a SynAmps RT 64-channel headbox and amplifiers and CURRYscan7 neuroimaging suite (Compumedics Neuroscan) at a sampling rate of 2 kHz. Eight electrodes were placed at Cz, CPz, C3, C4, Oz, FCz, T3, and T4. The ground electrode was placed at FPz and the reference electrode at Fz. The infant’s scalp was cleaned with NuPrep gel (D.O. Weaver and Co., Aurora, USA) to achieve impedances of 5-10 kΩ. A conductive paste (Elefix EEG paste, Nihon Kohden, Tokyo, Japan) was used to attach disposable Ag/AgCl cup EEG electrodes (Ambu Neuroline, Ballerup, Denmark) to the infant’s scalp. EEG was recorded for an average of 1.7 ± 0.7 hours (mean ± SD). ECG was co-registered with our EEG recordings using a single electrode (Ambu Neuroline, Ballerup, Denmark) placed on the infant’s chest (referenced to Fz).

To synchronise the vital signs and EEG recordings, we registered approximately 30 annotations simultaneously on both recordings using a bespoke triggering device (the PiNe box)^34^. Annotations were used to mark resting state as well as stimulus-evoked activity (these responses were used in other studies to investigate infant brain maturity – see Zandvoort *et al.* ^33^ – but were used here only to synchronise the EEG and vital signs recordings). The PiNe box enables synchronisation with an average precision of 105 ± 14 ms (mean ± SD)^34^. Annotation timings were matched between both vital signs and EEG recordings. Next, to further improve synchronisation, we utilised the ECG signals which were recorded on both the vital signs and EEG recording devices. To do so, ECG epochs were created from -10 to 10 seconds around each annotation. We aligned the ECG signals around each of these annotation epochs using cross correlations, providing a temporal delay between vital signs and EEG^34^. Temporal synchronisation between vital signs and EEG recordings was achieved by taking the median of the time differences across all annotations.

From the total of 123 recordings, we excluded 19 recordings due to (a) missing IP recordings (5 recordings), (b) missing annotations (9 recordings), and (c) inaccuracies in EEG-IP syncing (5 recordings; standard deviations over annotations exceeding 1 sec for annotation alignment or 0.1 sec during ECG alignment). Hence, data comprising 104 recordings from 68 infants were used in the analysis.

### Breath and apnoea identification

IP signals were processed to identify breaths (including apnoeas). To this end, we used an algorithm validated to detect inter-breath intervals in infants^14^. Briefly, the algorithm attenuates noise introduced by movements and cardiac activity by filtering the signal. An adaptive amplitude threshold then identifies individual breaths. This threshold was set at 0.4 times the standard deviation of the signal over the preceding 15 breaths. Finally, a support vector machine algorithm was applied to all long (here, 15 seconds or greater) inter-breath intervals to separate episodes of apnoea (cessation of respiration) and periods of low amplitude erroneously identified as apnoeas due to noise or shallow breathing. The inter-breath intervals marked as noise/shallow breathing were discarded from further analysis. The model was trained and validated on IP signals acquired during a clinical trial^35^, which was a completely independent data sample to the one used in the current study.

### Phase-amplitude coupling

In line with Osipova *et al.* ^12^, we defined phase-amplitude coupling (PAC) as cross-frequency coherence between a low-frequency phase and high-frequency amplitude. Note that the term “phase-amplitude coupling” is used interchangeably in the literature to mean the coupling principle of the statistical relationship between phases and amplitudes as well as the actual measure to detect this. We reserve the abbreviation for the latter. For our PAC measure, this meant that coherence was computed between the filtered IP signal and instantaneous amplitude of the EEG signal at some higher frequency (i.e., the second-order moment between the IP phase and EEG amplitude). More formally, in our case PAC was defined as:

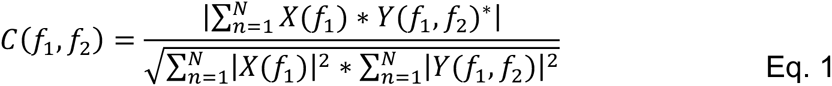

with *X* being the IP signal at frequency *f*_1_, *Y* the instantaneous EEG amplitude, and ^∗^ the complex conjugate. This instantaneous amplitude was obtained using the magnitude of the Hilbert transform after narrowly filtering the data at frequency *f*_2_. This component is often filtered with a bandwidth that relates to the low-frequency component *f*_1_. Following earlier recommendations, the filter bandwidth of the high-frequency component (at *f*_2_) was equal to a single time the frequency of the low-frequency component *f*_1_^36^ (e.g., when *f*_1_ equals 5 Hz the filter bandwidth of *f*_2_ is set to 5 Hz).

Data were segmented into 4-sec epochs (windows) (50% overlap), which meant that the frequency resolution was equal to 0.25 Hz (allowing to capture low delta-band frequencies), which comprised around four breaths. To focus on regular respiration, we discarded epochs which contained fewer than three detected breaths. An epoch was also removed from further analysis when its standard deviation exceeded 3 times the mean or median of the standard deviation of all epochs (which was done separately for the EEG and IP). Finally, we rejected any epoch containing an absolute value greater than 10 times the mean or median of the standard deviation of all epochs. Before cross-spectrum computation, EEG and IP epochs were convolved with a wavelet constructed from a Hanning window.

### Changes in PAC throughout the respiratory cycle

To investigate how cortico-respiratory coupling changes throughout the respiratory cycle, we estimated PAC at 50 equally spaced timepoints within the respiratory cycle. To this end, we time-locked both EEG and IP to each inter-breath interval as identified by the algorithm of Adjei *et al.* ^14^. At each respiratory cycle phase, cross- and auto-spectra were obtained to estimate PAC over all breaths within a recording. We limited the analysis to EEG channels FCz and Cz, because they exhibited the highest PAC magnitude (Figure 2), as well as statistically significant frequency-frequency samples relative to the surrogate PAC (see Figure 2D for the *p*-values of this statistical comparison). PAC was computed within the different frequency bands (delta: 0.5-4 Hz; and theta: 4-8 Hz) to see whether its modulation was specific for each frequency band. The same preprocessing steps apply as for PAC computation described earlier apart from the fact that data were segmented into 2/3 sec windows (so that temporal changes would appear within the respiratory cycle).

### Directionality of phase-amplitude coupling

To assess whether the cortical activity was preceding or trailing relative to the respiratory activity, we employed a cross-frequency version of the phase-slope index. Briefly, this metric estimates the phase lags over narrow frequency ranges and quantifies whether they increase or decrease over this range by estimating the phase-frequency slopes. Increasing or decreasing slopes signify changing time lags depending on the frequency of the oscillation and can therefore be indicative of whether a signal leads or trails another signal. We used the phase-slope index (Ψ) as proposed by Jiang *et al.* ^13^, which was defined as:

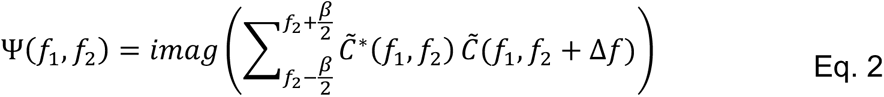

with *C̃* being the coherency (i.e., the normalised cross-spectrum) of *C* defined in Eq. 1. Δ*f* is the frequency resolution and *β* the bandwidth used to estimate the phase slope. In line with previous recommendations by Jiang *et al.* ^13^, we set *β* to 1 Hz (i.e., four times the frequency resolution).

### Surrogate analysis

To assess the statistical significance of PAC (i.e., coherence) and phase-slope index at each of the EEG channels, we performed a surrogate analysis under the null hypothesis that the means of the true and surrogate PAC (and phase-slope index) were equal. The surrogate analysis was performed by randomly permuting the epoch (4-s windows used for PAC calculation – see previous sections) order of the EEG amplitude while leaving the respiration epoch order unchanged (i.e., respiration segment/epoch S1 was paired with an EEG segment/epoch Sj, j ≠ 1). Importantly, no temporal samples were shuffled within epochs. Thus, the full within-epoch temporal structure, including autocorrelation and spectral profile (auto-spectra), was preserved for both signals. This permutation destroys trial-wise cross-signal phase alignment (and therefore coherence) while retaining the intrinsic dynamics of each signal. We obtained a surrogate spectrum for each recording. Next, the true and surrogate spectra were statistically compared on a sample-by-sample basis using dependent *t*-tests^37^ (using the FieldTrip toolbox – version 20210325). This comparison was performed over all recordings. Statistical significance was defined using the false discovery rate correction (with an alpha level set at 0.05 [phase-slope index] or 0.001 [coherence]).

### Statistical analysis of relationships between cortico-respiratory coupling and apnoea

We tested our main hypothesis that PAC is negatively associated with apnoea rate. Vital signs were continuously monitored for the infants that took part in the *Breathing and Brain Development* study, which allowed us to analyse their vital signs in longer time windows than the EEG. We here selected up to 24 hours of vital signs from the day of the EEG recording (mean ± standard deviation: 9.5 ± 9.9 hours) and expressed this as a rate per hour. We then constructed linear mixed-effects models to study the relationships between PAC and apnoea rate. We included the coherence magnitude as a fixed factor, which was averaged over the statistically significant samples from channels FCz and Cz (see Figure S4 for the channel-specific relationships). As fixed effects, we included (1) data length of the respiratory signal, (2) mode of ventilation (‘Self-ventilating’, ‘Low flow’, ‘High flow’; Figure S8), and (3) PMA (Figure S7). Infant ID was included as a random effect acting on the intercept. Data length, mode of ventilation and PMA were included as confounding factors. P-values (for the regression coefficient of PAC) and partial correlation coefficients were reported from these models. The normality assumption was verified with Q-Q plots. We also tested the linearity assumption (see Supplementary text). For visualisation (Figure 4), we used the plotAdjustedResponse function in MATLAB to adjust for the fixed factors (calculating a separate model without the random effect of an infant). Regression lines are visualised by calculating the line-of-best-fit on the adjusted data.

