## Supplementary material for "Cortical motor activity modulates respiration and reduces apnoea in neonates"

### ***Supplementary text***

*Assessing the linearity of factors*

Using multivariable fractional polynomial models, we tested the assumption of linearity for the linear mixed-effects model (Figure 4A). Polynomial models determine whether relationships are better modelled through non-linear relationships by finding the most appropriate functional form for continuous predictors. Using the *mfp* package in Rstudio (Version 2024.04.1+748), we estimated these forms and statistically tested them against the factor being modelled linearly. The same factors were included as presented in the main text (Figure 4A; i.e., mode of ventilation, post-menstrual age, data length when estimating apnoea rate and infant). This resulted in a *p*-value of 0.74, confirming the linearity assumption. The *p*-value represents the difference in deviances between the fractional polynomial and linear models.

### *Supplementary figures*

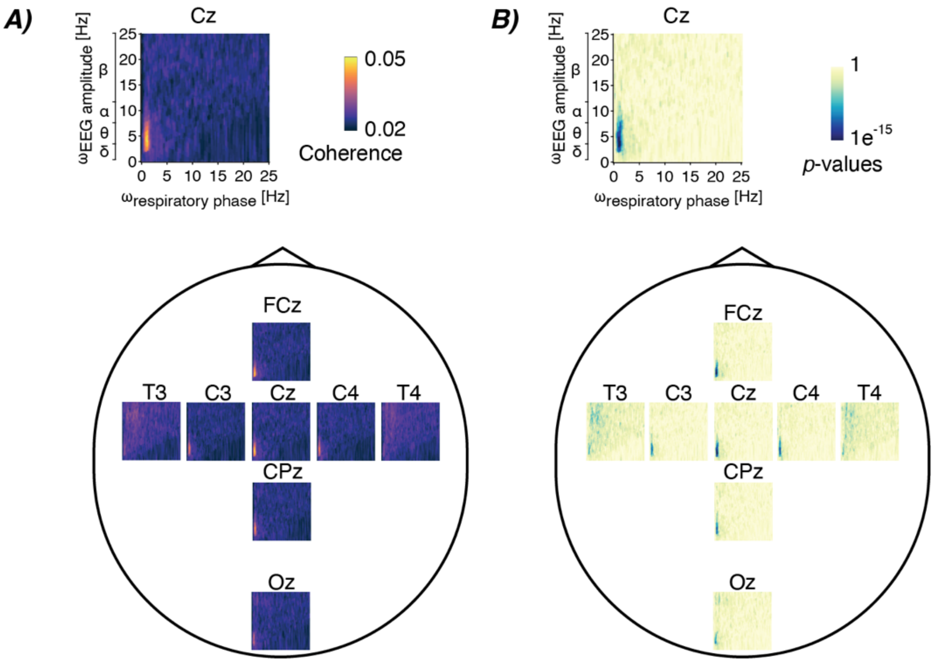


***Figure S1. Cortico-respiratory coupling in infants**. **A)** Spatial map of coherence-based phase-amplitude coupling between respiratory phase ($\omega_{respiratory\ phase}$) and EEG amplitude ($\omega_{EEG\ amplitude}$) over full frequency ranges (Figure 2 depicts the zoomed-in spectra). **B)** Analogous spatial map for the (uncorrected) statistical significance relative to surrogate PAC (reported on a logarithmic scale).*

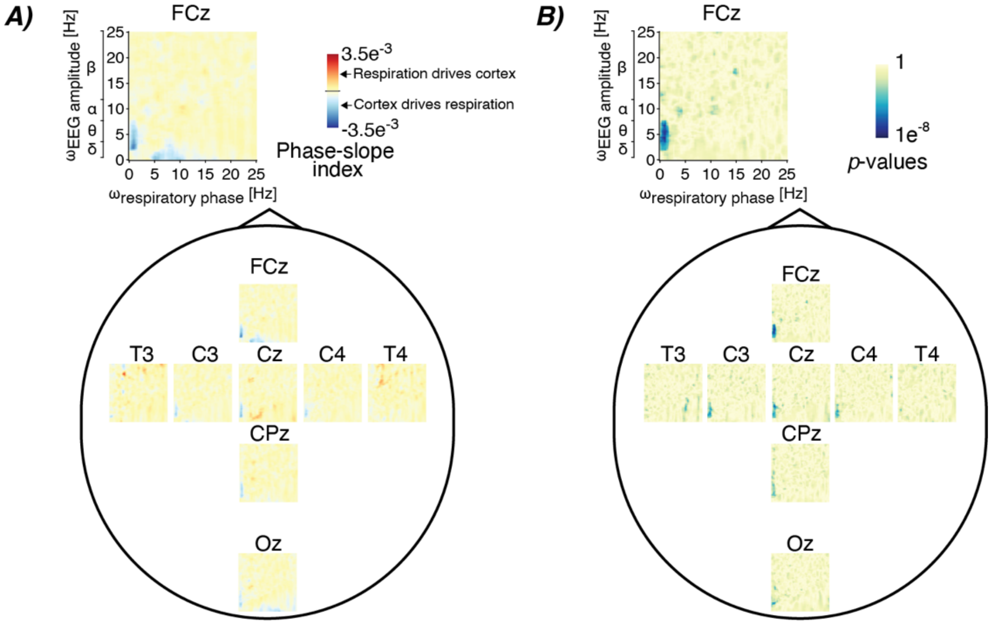


***Figure S2. Directionality of the cortico-respiratory coupling**. **A)** Spatial map of the phase-slope index between the respiratory phase ($\omega_{respiratory\ phase}$) and EEG amplitude ($\omega_{EEG\ amplitude}$), here shown over full frequency ranges (see Figure 3 for the zoomed-in spectra). **B)** The corresponding (uncorrected) statistical significance (reported on a logarithmic scale).*

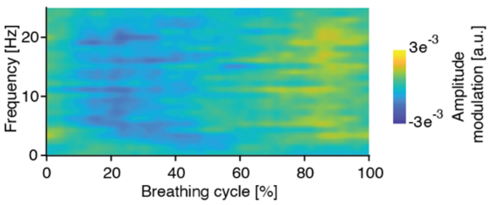


***Figure S3. Photoplethysmographic amplitude modulation depending on the respiratory cycle and frequency.***

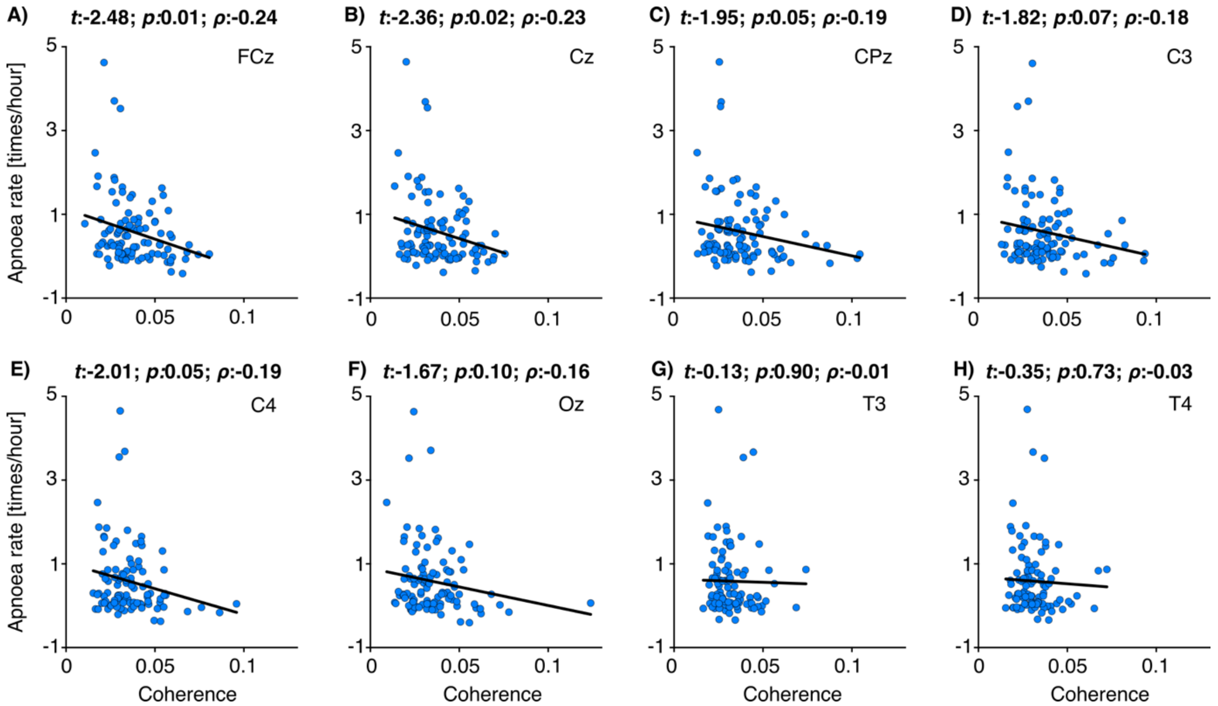


***Figure S4. Relationships between phase-amplitude coupling and apnoea rate for individual EEG channels.*** *Each blue dot is the data of an individual recording (data of 68 infants were included from 104 test occasions). Black linear graphs are the best fit of the linear mixed-effects models (fixed factors: coupling averaged over statistically significant samples at each electrode respectively, data length to determine apnoea rate, mode of ventilation, and postmenstrual age; random factor: infant [random intercept]). Panel titles contain the test statistic of the predictor's regression slope (t), its significance (p), and partial correlation coefficient (ρ). Electrode position is indicated in top right of each panel, with* ***A)*** *FCz,* ***B)*** *Cz,* ***C)*** *CPz,* ***D)*** *C3,* ***E)*** *C4,* ***F)*** *Oz,* ***G)*** *T3, and* ***H)*** *T4. Electrodes with weaker coherence overall (see Figure 2C,D) – Oz, T3, T4 – did not show a significant relationship between apnoea rate and coherence as expected as these electrodes do not overlay cortical motor areas.*

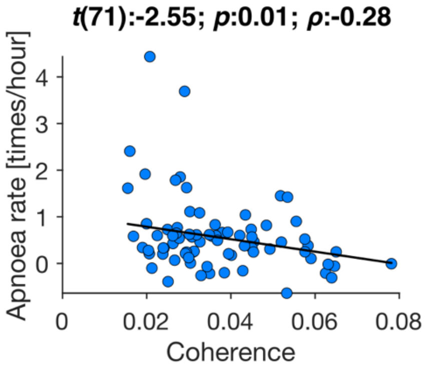


***Figure S5. Apnoea rate for preterm infants compared with phase-amplitude coupling for preterm infants.*** *Relationship between PAC (defined as cross-frequency coherence) and apnoea rate for the preterm infants (≤36 weeks postmenstrual age) (data of all infants is presented in Figure 4). Each blue dot is the data of an individual recording (data of 53 infants were included from 86 test occasions). Black linear graph is the best fit of the linear mixed-effects model (fixed factors: coupling averaged over statistically significant samples from channels FCz and Cz (see Figure S4 for channel-specific relationships), data length to determine apnoea rate, mode of ventilation, and postmenstrual age; random factor: infant [random intercept]). Panel title contains the test statistic of the predictor's regression slope (t), its significance (p), and partial correlation coefficient (ρ).*

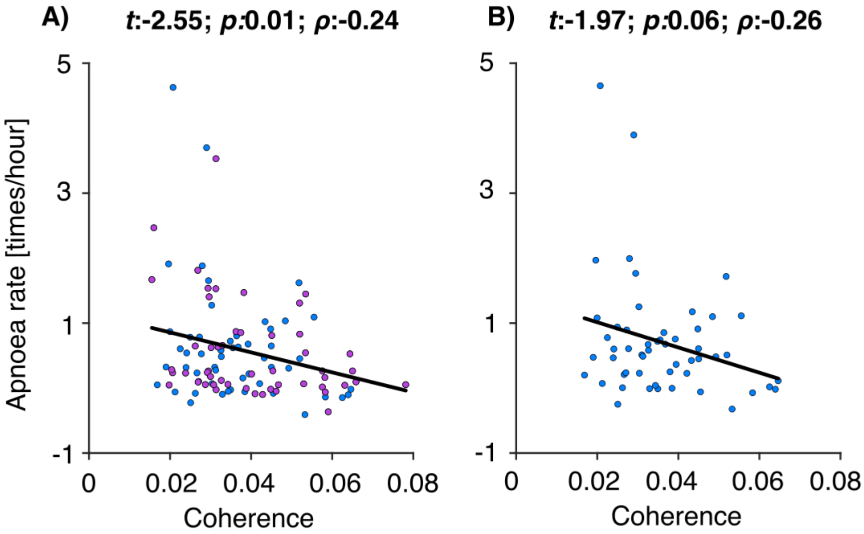


***Figure S6. Relationship between apnoea rate and phase-amplitude coupling in infants receiving caffeine. A)*** *Data for all recordings, with blue dots indicating sessions where infants had received caffeine (medically prescribed), and purple dots indicate sessions where infants did not have caffeine prescribed on the day of recording (total: 68 infants on 104 recording sessions).* ***B)*** *Stratified analysis for infants receiving caffeine (35 infants on 53 recording sessions). Infants receiving caffeine are more likely to experience apnoeas (which is why caffeine is prescribed). Each dot is the data of an individual recording. Black linear graph is the best fit of the linear mixed-effects model (fixed factors: coupling averaged over statistically significant samples from channels FCz and Cz, data length to determine apnoea rate, mode of ventilation, and postmenstrual age; random factor: infant [random intercept]). Panel title contains the test statistic of the predictor's regression slope (t), its significance (p), and partial correlation coefficient ($\rho$).*

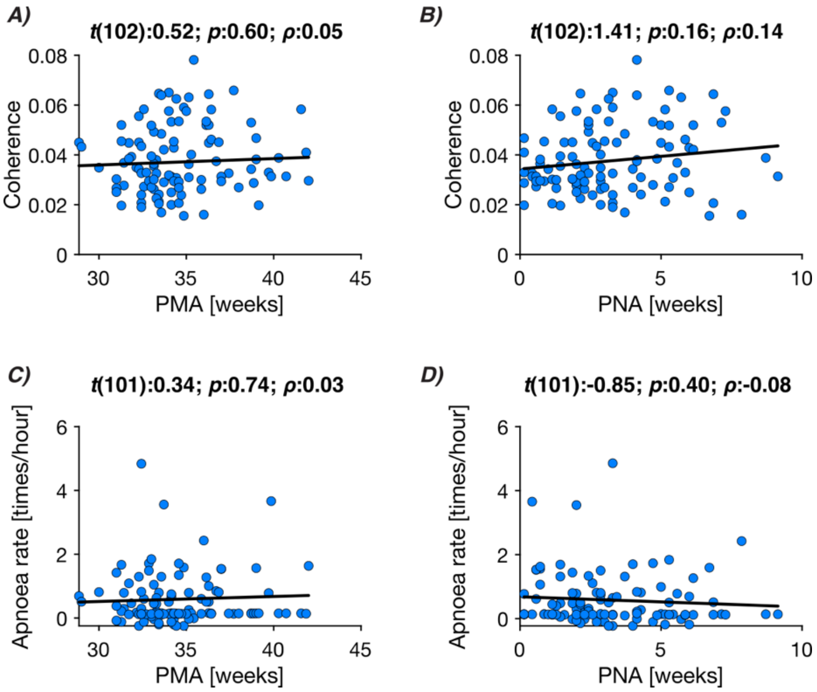


***Figure S7. Coupling and apnoea rate relationships with age. A)*** *Phase-amplitude coupling (defined as cross-frequency coherence), and* ***B)*** *apnoea rate relationships with postmenstrual age (PMA).* ***C-D)*** *Same relationships over postnatal age (PNA). Each dot is the data of an individual recording (data from 68 infants on 104 test occasions were included). Black linear graphs are the best fit of the linear mixed-effects models (fixed factor: postmenstrual age or postnatal age; random factor: infant – in line with previous publications, the data length of the respiratory signal was additionally included in the models of panels B and D). Panel titles describe the predictor's test statistic of the regression slope (t), its significance (p), and partial correlation coefficient ($\rho$).*

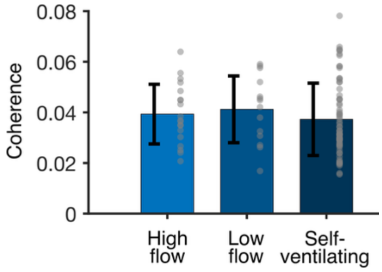


***Figure S8. Cortico-respiratory coupling as a function of mode of ventilation.*** *Coherence-based PAC were averaged for each recording using the statistical masking obtained from Figure 2.*
